# Auditory enhancement of illusory contour perception

**DOI:** 10.1101/860965

**Authors:** Ruxandra I. Tivadar, Anna Gaglianese, Micah M. Murray

## Abstract

Illusory contours (ICs) are borders that are perceived in the absence of contrast gradients. Until recently, IC processes were considered exclusively visual in nature and presumed to be unaffected by information from other senses. Electrophysiological data in humans indicates that sounds can enhance IC processes. Despite cross-modal enhancement being observed at the neurophysiological level, to date there is no evidence of direct amplification of behavioural performance in IC processing by sounds. We addressed this knowledge gap. Healthy adults (N=15) discriminated instances when inducers were arranged to form an IC from instances when no IC was formed (NC). Inducers were low-constrast and masked, and there was continuous background acoustic noise throughout a block of trials. On half of the trials, i.e. independently of IC vs. NC, a 1000Hz tone was presented synchronously with the inducer stimuli. Sound presence improved the accuracy of indicating when an IC was presented, but had no impact on performance with NC stimuli (significant IC presence/absence × Sound presence/absence interaction). There was no evidence that this was due to general alerting or to a speed-accuracy trade-off (no main effect of sound presence on accuracy rates and no comparable significant interaction on reaction times). Moreover, sound presence increased sensitivity and reduced bias on the IC vs. NC discrimination task. These results demonstrate that multisensory processes augment mid-level visual functions, exemplified by IC processes. Aside from the impact on neurobiological and computational models of vision our findings may prove clinically beneficial for low-vision or sight-restored patients.

## Introduction

Everyday vision must oftentimes overcome impoverished viewing conditions for the correct perception of our environment. These conditions can include poor lighting, occlusion, etc. Our brains must infer the information that our visual system does not see, including borders and shapes across regions of isoluminance or when contrast gradients are missing (Lesher, 1995). The perception of such illusory contours (ICs) is an evolutionarily conserved process and has been reported in animals including zebrafish, frogs, owls, cats, dogs and monkeys (among others) (reviewed in Murray & Herrmann, 2013). Illusory contour perception is classically considered a purely visual function that occurs independently of processing of other sensory information.

Results of research from the past ~20 years have demonstrated that multisensory processes are pervasive and impact perception and behaviour even at the earliest cerebral stages (reviewed in Murray, Lewkowicz, Amedi, & Wallace, 2016). For example, auditory information can impact the excitability of primary visual cortex, can facilitate stimulus detection, and can even implicitly facilitate memory encoding and retrieval (reviewed in Matusz, Wallace, & Murray, 2017; Murray, Thelen, et al., 2016). Auditory information can likewise impact visual attention processes (e.g. Matusz & Eimer, 2011; Spence, 2010), visual discrimination and object recognition (e.g. Amedi, von Kriegstein, van Atteveldt, Beauchamp, & Naumer, 2005), as well as visual recognition memory (reviewed in Matusz et al., 2017). All of these (and other) examples entail auditory influences on responses to or representations of physically-present visual information. By contrast, it remains largely unknown whether multisensory processes would impact the processing of ICs at a behavioural level. Given recent evidence from our group indicating that neural mechanisms of IC perception can be impacted by task-irrelevant sounds (Tivadar, Retsa, Turoman, Matusz, & Murray, 2018), we here wanted to investigate behavioural effects of this enhancement. In the present study, we therefore presented healthy, sighted individuals with low contrast IC stimuli (and their controls – i.e. No Contour stimuli, NCs) in a visual masking paradigm (Ringach & Shapley, 1996) to avoid a ceiling effect on performance. Although sounds were presented on only half of the trials, participants performed the experiment in a setting of continuous background acoustic noise (i.e. the sound of an MRI scanner). We hypothesized that sounds would facilitate IC sensitivity without a concomitant a general alerting or arousal effect.

## Methods

### Participants

We tested 18 right-handed, neurologically healthy participants (10 women; mean±SD age: 27.7±4.0 years). All participants provided written, informed consent to procedures approved by the Cantonal Ethics Committee (protocol 2018-00240). All participants but one were right-handed (Oldfield, 1971), all had normal or corrected-to-normal vision, and all reported normal hearing capacities. None reported any history of psychiatric or neurological illness. Data from one participant were excluded due to failure to comply with the task instructions (i.e. a preponderance of anticipatory button presses), and data from another two participants were excluded due to ceiling level performance and post-experiment debriefing that revealed these individuals had extensive low-contrast visual expertise. Thus, our final dataset comprised 15 participants (9 women; mean±SD age: 27.1±3.7 years). A power analysis using G*Power (version 3.1.9.2) (Faul, Erdfelder, Lang, & Buchner, 2007) indicated that a sample size of 15 was necessary to obtain a statistical power of at least 0.95 for detecting an effect of a moderate size (f = 0.40) with our 2×2 factorial design and assuming a correlation of 0.50 between repeated measures. We also chose this sample size because our previous studies have obtained highly consistent data with similar sample sizes (Tivadar et al., 2018).

### Stimuli and task

Stimuli were comprised of a set of 4 circular Kanizsa-type (Kanizsa,1976) ‘pacmen’ inducers (i.e. a circle with a rectangular ‘mouth’ missing) that were arranged to either form an illusory line or not (hereafter IC and NC conditions, respectively) (see **Figure 1A**). Each inducer subtended 2° in diameter of visual angle at a viewing distance of 80cm from fixation, which was measured before the beginning of the experiment. On a given IC trial, the four pacmen were positioned along the upper and lower horizontal and left and right vertical axes to form a single rectangular IC in the upper, lower, left, or right visual field with the nearest “illusory” edge at 2.48° eccentricity from central fixation. For NC configurations, each pacman was rotated 90° towards the next one. Pacmen stimuli always subtended 3.5° centre-to-centre eccentricity, with a support ratio of = 0.4, (i.e. the ratio of physically present versus illusory borders of the rectangle). There were two different variations of the NC displays, which were automatically randomised within a block of trials. These variations in how ICs were created were included to prevent participants from selectively attending to particular regions of space or to a specific pacman inducer as a strategy to successfully complete the task. Inducers were black on a dark grey background (RGB values of 0, 0, 0 and 6, 6, 6, respectively). The employed forms, created from the aligned orientation of 2 of the 4 inducers, have been used in prior IC studies by our group, and are known to result in robust IC sensitivity (Anken, Tivadar, Knebel, & Murray, 2018). We would emphasize that the orientation of any single inducer was therefore uninformative as to whether or not the stimulus contained an IC. In both IC and NC conditions, synchronous to the presentation of the visual stimuli, an auditory stimulus was presented on half of the trials. The auditory stimulus was generated with Audacity freeware (available at http://www.audacityteam.org). It was a 1000Hz sinusoidal pure tone of 100ms duration with 10ms fade in/out to avoid clicks. The sample rate was 44.1kHz and the stimulus was presented in 16 bits per sample. For the no-sound condition, a silent sound was generated in Audacity, lasting for 100msec. The generation of this silent sound was done in order to ensure the uniform distribution of sound and no-sound conditions. Sounds were presented via in-built 13-inch Mac loudspeakers in a sound attenuated room (WhisperRoom MDL 102126E), and the sound volume was kept at a uniform level (79.3dB as measured at the distance of the head using a CESVA SC-L sound pressure meter). Moreover, a 1-minute recording of a 7T Siemens MRI scanner noise was added to each block. The reasoning was twofold. On the one hand, we wanted to avoid any potential alerting effect of the task-irrelevant sounds. On the other hand, we anticipate conducting future work during the acquisition of MRI data. Each block contained 300 visual stimuli with unequal probability of IC (i.e. 120 trials, 30 trials per IC location) and NC conditions (i.e. 30 trials), but equal probability of sound or no sound conditions (150 trials each). Each participant completed three blocks, in which all the conditions were randomly presented. The stimulus sequence is schematized in **Figure 1B**. The subjects sat in a sound-attenuated dark room, at a distance of 80cm from the presentation screen. Stimuli were presented for 100ms with an inter-stimulus interval ranging between 500ms and 1000ms. A mask, consisting of filled disks (with the same radius, position, and contrast as the inducers) as well as a black central fixation dot were presented on a dark grey background, and remained on the computer screen for the whole duration of the experiment, excepting the 100ms when the IC/NC stimulus appeared. Pilot studies helped to identify low contrast inducer stimuli as well as a masking procedure that together resulted in task performance below ceiling levels. The participants’ task was a two-alternative forced choice that required the discrimination between IC and NC presence on each trial via a right-handed button press. Stimulus delivery and behavioural response collection were controlled by Psychopy software (Peirce, 2007). During the experiment, participants took regular breaks between blocks of trials to maintain high concentration and prevent fatigue.

**Figure 1.**
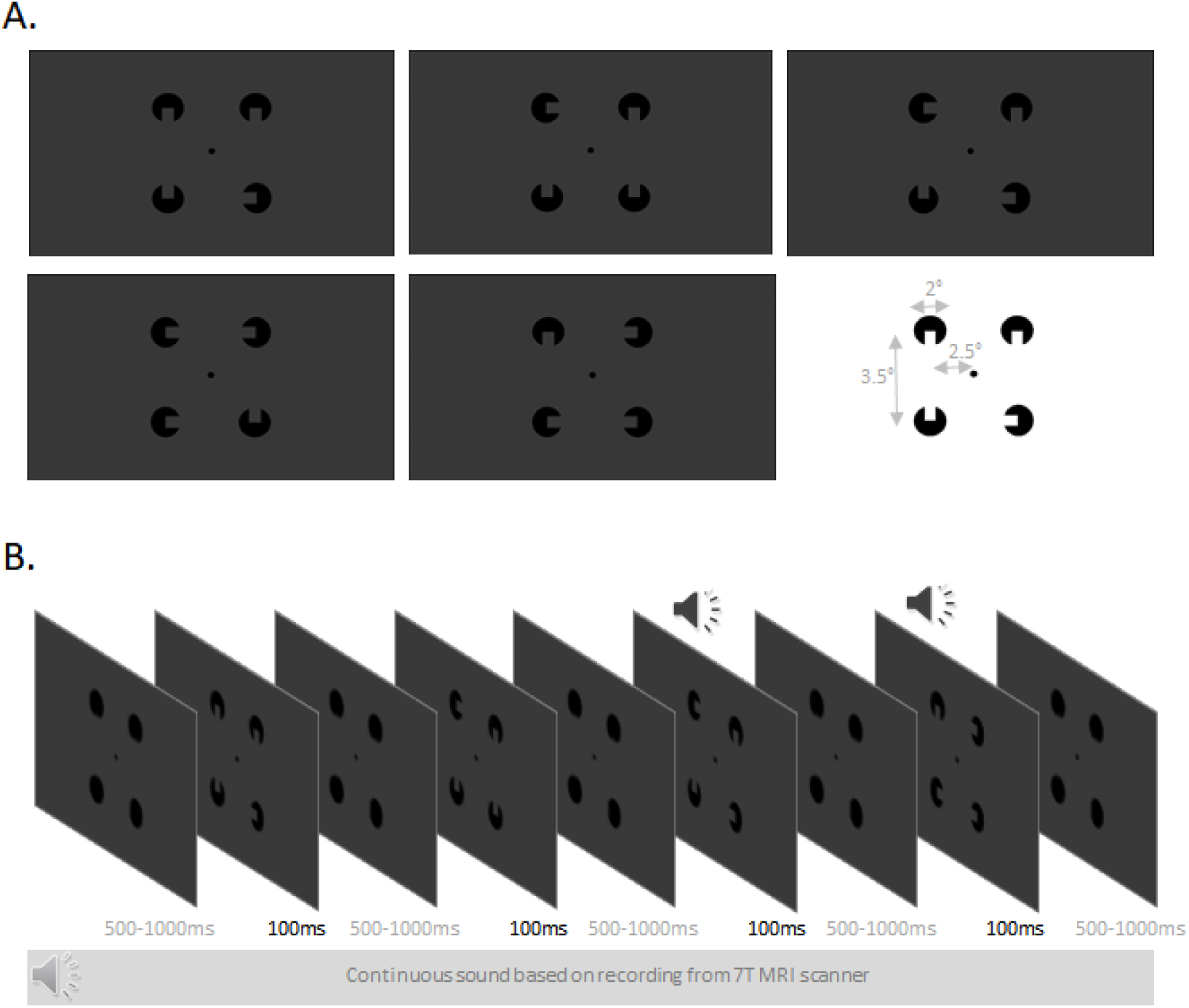
Stimuli and schematic of trial structure. **A.** Examples of stimulus configurations show how IC conditions resulted in the perception of a line on the left, right, above, or below central fixation. In the NC condition no line was perceived. Note that stimulus contrast has been enhanced for illustration purposes. The inset indicates the size and eccentricity of the pacmen inducers and IC line. **B.** Masking stimuli were the four filled circular inducers that were presented for a duration varying between 500ms and 1000ms. The task-relevant stimuli were then presented for 100ms on any given trial, and participants were instructed to indicate whether or not an illusory contour had been presented. On half of the trials, a task-irrelevant 1000Hz pure tone was also presented in synchrony with the visual stimulus. During a block of trials there was the continuous presentation of a background sound, which was the recording of the 7T Siemens scanner noise. The reader should note that the orientation of any individual inducer was in and of itself uninformative of whether or not an illusory contour was present or absent.

### Behavioural Analyses

Our general approach was to first test if each dependent measure was normally distributed. If so, then parametric statistical tests were used. If not, then non-parametric tests were used. The percent correct data did not fulfil the assumptions of normality and homoscedasticity, which were tested via Shapiro-Wilk and Levene tests, respectively. Therefore, we ran a permutation-based 2×2 multivariate ANOVA (Wheeler & Torchiano, 2010) with factors Sound presence/absence and IC presence/absence (IC/NC). The random seed value to iniate the permuations was set and fixed at 42, in order to obtain replicable results. Reaction time (RT) data were normally disributed (Shapiro-Wilk test). RTs were therefore tested with a parametric repeated measures analysis of variance (rmANOVA), following a 2×2 within subjects factorial design (IC presence/absence × sound presence/absence). Only correct trials were used for the RT analysis. We first excluded outlier trials on a single subject basis (i.e. for each subject and condition), applying a mean ±2 standard deviations criterion (Ratcliff, 1993). On average, 2.7% of the trials were excluded from any condition.

We also computed the sensitivity and bias of the IC versus NC discrimination both for when sounds were absent and present, using the measures A and beta, since our data were not normally distributed as detailed above (Zhang & Mueller, 2005). Hits were trials when ICs were present and the participant reported an IC. Misses were trials when ICs were present and the participant reported an NC. Correct rejections were trials when NCs were presented and the participant reported an NC. False alarms were trials when NCs were presented and the participant reported an IC. This measure of sensitivity (A) can range from 0 to 1, with larger values indicative of higher discriminability. Likewise, this measure of decision bias (b) can range from 0 to 1, with values closer to 1 indicative of a bias for a ‘yes’ response and values closer to 0 indicative of bias for a ‘no’ response. Given our hypothesis that sounds would increase sensitivity and decrease bias, we used 1-tailed Wilcoxon ranked sum tests.

## Results

**Figure 2** displays the group-average percentage correct and RTs for each condition, and the insets display the difference in percentage correct for the IC and NC conditions when the sound was absent versus present for each individual participant. The permutation tests on the percent correct data showed a significant main effect of IC presence/absence (*p*<0.01, number of iterations: 5000), as well as a Sound × IC presence/absence interaction (*p*=0.02, number of iterations: 5000). The main effect of Sound was not significant (*p*= 0.06, number of iterations: 5000). We then ran a series of Wilcoxon signed rank tests to understand the basis for the significant interaction. There was a significant difference in percentage correct with IC stimuli when the sound was present vs. absent (81.5% vs. 77.7%; p=0.012). By contrast, there was no such difference with NC stimuli (44.7% vs. 45.4%; p=0.410). The 2×2 rmANOVA on RTs revealed a main effect of IC presence/absence (F_(1,14)_ =10.0; p=0.007; η_p_^2^=0.42), that was due to generally faster RTs for IC than for NC stimuli (753ms vs. 956ms). There was also a main effect of Sound presence/absence (F_(1,14)_ =47.2; p<0.001; η_p_^2^=0.77) that was due to generally faster RTs when sounds were present than absent (834ms vs. 874ms). However, there was no reliable interaction between IC presence/absence and Sound presence/absence (F_(1,14)_<1; p=0.55; η_p_^2^=0.03). The 1-tailed Wilcoxon signed rank tests on sensitivity and bias confirmed our hypothesis that sound presence increased sensitivity (0.695 vs. 0.679; p=0.038) and decreased bias (0.650 vs. 0.683; p=0.02). **Figure 3** displays normalized changes, expressed as a percent, between sound absent and present conditions for sensitivity and bias for each participant.

**Figure 2.**
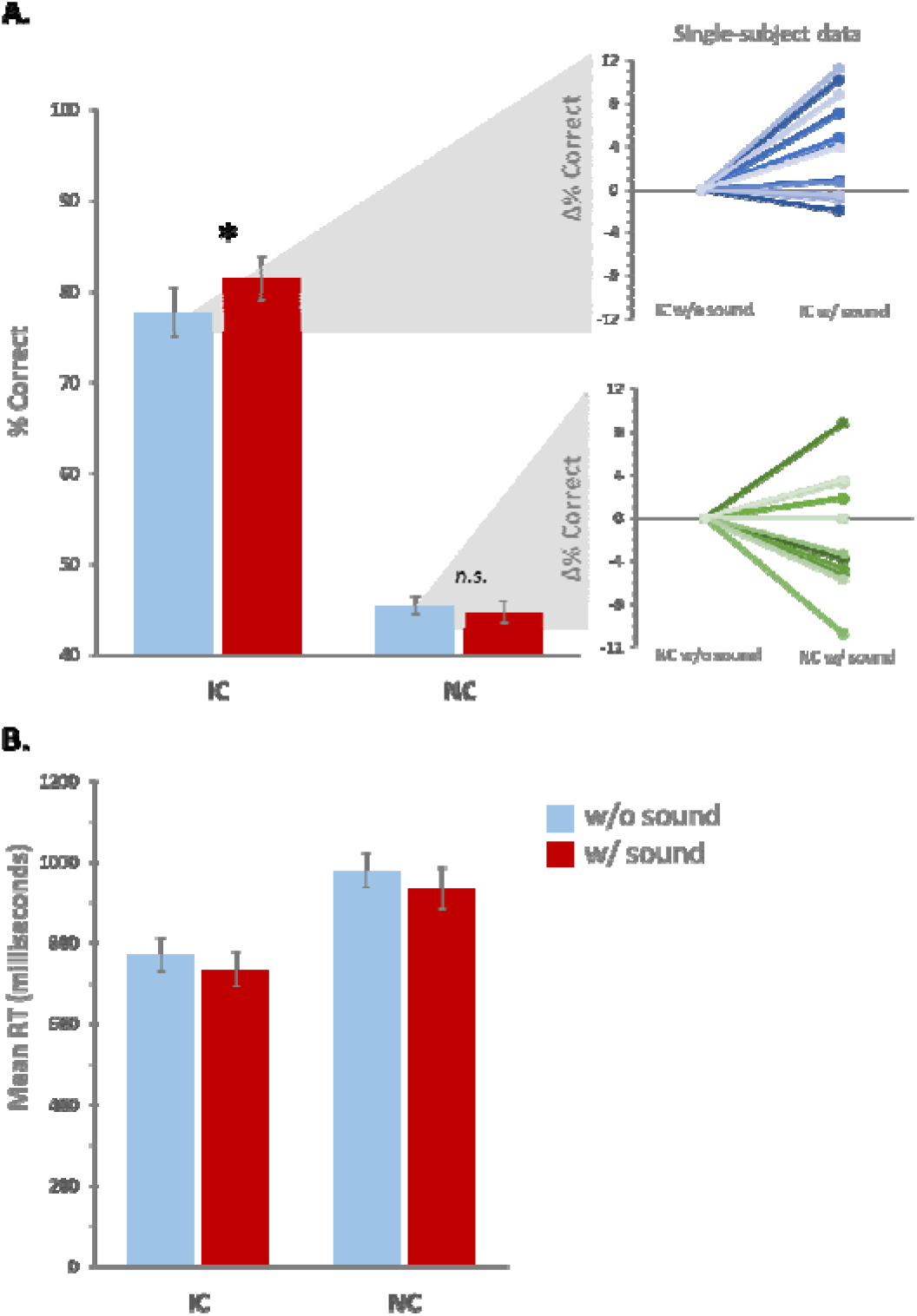
Group-averaged (N=15) percent correct (A) and reaction time (B) for each of the experimental conditions. Error bars indicate the standard error of the mean. The insets display single-subject data for the difference between IC conditions and between NC conditions as a function of sound presence (normalized to the condition without sounds).

**Figure 3.**
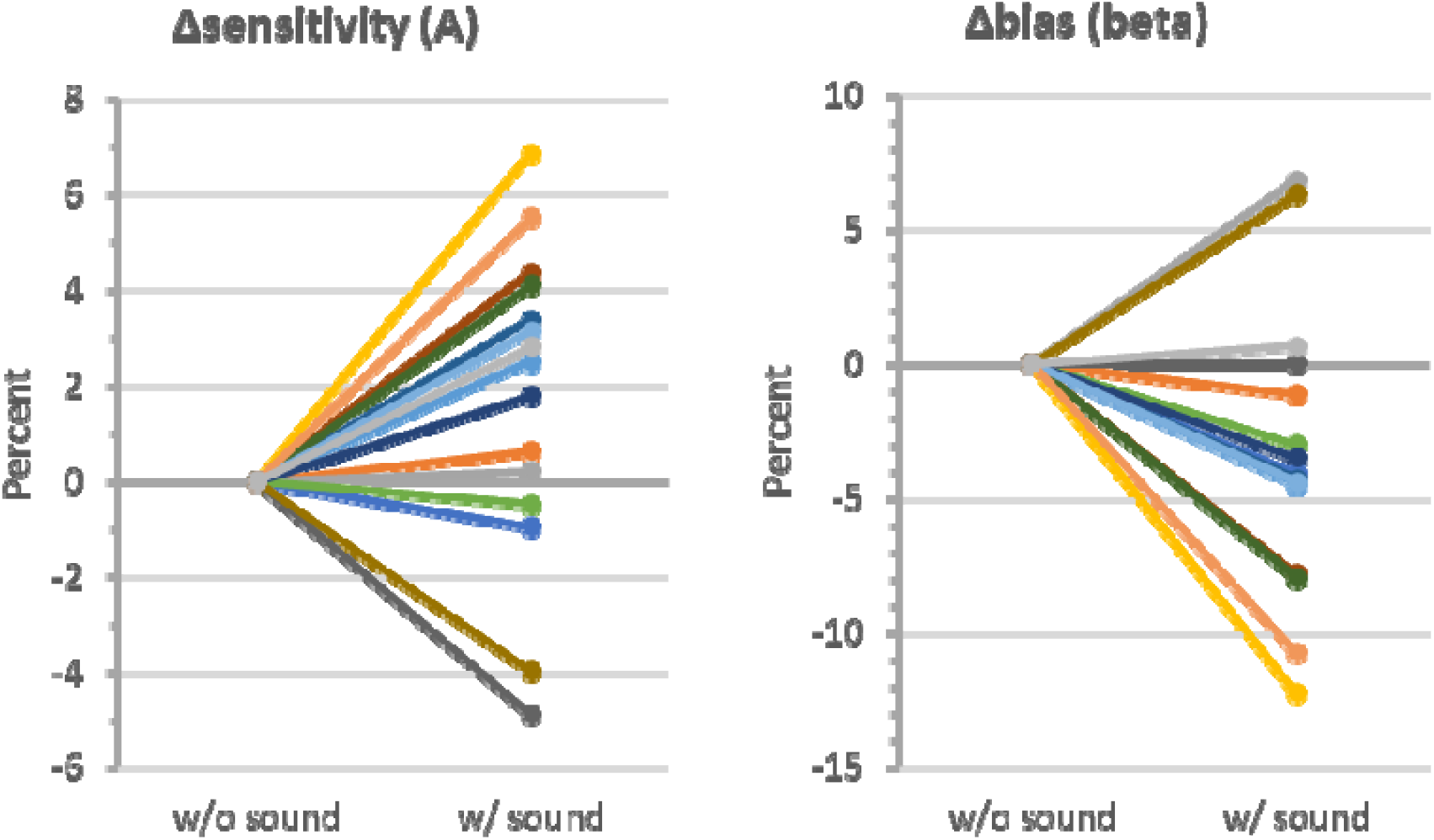
Single-subject data for the percent difference in sensitivity and percent difference in bias as a function of sound presence (normalized to the condition without sounds). Colours are the same across the graphs for a given participant. Sensitivity increased while bias decreased with sound presence.

## Discussion

Based on our prior electroencephalographic findings (Tivadar et al., 2018), we hypothesized that sounds would facilitate IC sensitivity without a concomitant a general alerting or arousal effect. Our speculation was that such was not observed in our prior work because performance was at near-ceiling levels. The present study therefore strived to make the task more challenging and thus increase the propensity for cross-modal effects. This was borne out in our results. The ability to report the presence of an illusory contour is facilitated by the synchronous presentation of a sound. Sounds did not affect performance on trials when no contour was present. This was the case despite the sound being completely uninformative as to whether or not a given trial indeed contained an illusory contour. Sounds were equally probable for both IC and NC conditions. Two aspects in our design and empirical findings help to further exclude an account based on potential arousal or alerting effects of sound or any effects due to the sudden temporal onset of an acoustic stimulus. First, we included a continuous background auditory stimulus (i.e. the recording of an MRI scanner acquisition). Second, while sounds increased our measure of sensitivity, there was a concomitant decrease in bias. Thus, what was hitherto considered an exclusively visual process is instead also subject to cross-modal influences. Sounds impact visual completion processes, extending the breadth of multisensory phenomena to include mid-level vision and the construction of visual perception.

The fact that sounds affected accuracy on the IC, but not NC condition, provides evidence that sounds in our experiment were affecting the visual completion process and not (or not exclusively) the perception of the inducers themselves. For example, it might be posited that sounds enhance the perceived brightness of the physically-present inducer stimuli. Indeed, our and others’ prior works provide evidence for precisely such a phenomenon (reviewed in (Murray, Thelen, et al., 2016)). However, in such studies the task entailed stimulus detection, with visual stimulus presentation being the independent variable. Instead, each trial in the current experiment included a visual stimulus, with the presence versus absence of illusory contours being the independent variable. It may thus be the case that any influence of sounds depends critically on task contingencies; something prior studies would indeed suggest to be the case in general with regard to multisensory processes (reviewed in (ten Oever et al., 2016)) and perhaps also specifically in the case of illusory contour processing. For example, Fiebelkorn et al. (2010) failed to observed behavioural or electrophysiological enhancement of illusory contour processes by sounds in their task involving detection of a flickering inducer stimulus. Instead, there was a general reduction in hit rates under multisensory versus unisensory conditions. Tivadar et al. (2018) observed electrophysiological enhancement of event-related potential indices of illusory contour sensitivity despite near-ceiling performance and thus the absence of any concomitant behavioural effects when the task involved discrimination of IC presence versus absence.

The present results lend further support to the notion that sounds can enhance the excitability of visual cortices. In our prior work (Tivadar et al., 2018), we observed that an event-related potential signature of illusory contour sensitivity was enhanced by the presence of task-irrelevant sounds. A network of brain regions involved in this effect included not only LOC and the intra-parietal lobule, as previously observed in ERP studies of IC sensitivity (Murray & Herrmann, 2013), but also V1. The implication of V1 in IC processing has remained somewhat elusive in prior visual-only studies in humans (Murray & Herrmann, 2013). This network of brain regions was functionally correlated only under the multisensory condition, as compared to the visual-only condition. Crucially, amodal stimuli did not elicit such effects, which speaks against a direct effect of sounds on shape completion processes. Rather, our contention was that sounds affected the perceived brightness of illusory contour figures, possibly by enhancing the excitability of neurons in V1; a mechanism supported by transcranial magnetic stimulation studies of cross-modal influences on visual cortex excitability (e.g. (Romei, Murray, Cappe, & Thut, 2009, 2013; Romei, Murray, Merabet, & Thut, 2007; Spierer, Manuel, Bueti, & Murray, 2013). What these collective data show is that sounds, including 1000Hz tones used here, enhance the excitability of visual cortex over a time window spanning ~30-150ms after sound onset. These timings can be placed alongside work characterising the latency and spread of visually-driven inputs within visual cortices, where response onset in areas V1, V2, V4, STS, IP, and IT of the non-human primate all occurred over the ~30-70ms post-stimulus interval (e.g. (C E Schroeder, Mehta, & Givre, 1998). If one applies a 3:5 ratio (Musacchia and Schroeder, 2009 Hearing Research) to ‘convert’ these latencies from non-human primates to humans, one obtains a range of ~40-100ms. The implication is that sounds can, in principle, increase visual cortical excitability across a wide swath of cortical regions contemporaneously with or even before the arrival of the feedforward visually-driven input signal.

Phase-resetting of ongoing alpha activity by sounds (Romei, Gross, & Thut, 2012) may be a neurophysiologic mechanism contributing to these excitability increases (Ohshiro, Angelaki, & DeAngelis, 2017; van Atteveldt, Murray, Thut, & Schroeder, 2014). Specifically, LOC feedback input might modulate ambient oscillatory activity in V1 neurons, which may be in a more excitable state due to a modulatory effect of sounds. In fact, there are cells in V1 that exhibit bipolar properties, such as layer-3 complex pyramidal cells that excite one another monosynaptically via horizontal connections, and inhibit one another via disynaptic inhibition (Grossberg, Mingolla, & Ross, 1997). Modelling data would suggest that such cells fire only when their receptive fields lie between aligned inducers of ICs, but not when they lie beyond a single inducer. Sounds might, in turn, alter activity in such cells by aligning neuronal excitability to expected visual modulations, which foster the extraction of the most relevant visual cues, as is observed in the case of visual effects on ongoing auditory processing (Charles E Schroeder, Lakatos, Kajikawa, Partan, & Puce, 2008). Such cross-modally triggered phase locking of perceptually relevant oscillatory alpha activity is thought to be evoked either by the sounds themselves, or by phase-resetting of ongoing oscillations (Romei et al., 2012).

Our findings also provide an important proof-of-principle for the application of multisensory approaches as a particularly cost-effective means for low-vision rehabilitation. Results from early blind or early visually impaired individuals (for example, through congenital cataracts) emphasize the wide-ranging effects that partial or total visual deprivation can have on cognitive functioning (Murray, Matusz, & Amedi, 2015). For example, (McKyton, Ben-Zion, Doron, & Zohary, 2015) tested a sample of children suffering from congenital cataracts that were operated only at the age of about 9 to 11 years. They found intact low-level processing in this sight-restored population, with children performing almost as well as sighted counterparts on tasks that required colour, size or shape discrimination of visual stimuli (McKyton et al., 2015); a result also documented by previous research in a similar population (Maurer, Lewis, & Brent, 1989). Nevertheless, when the visual tasks required use of mid-level visual processing, such as discrimination of stimuli based on shading, illusory contour completion, or stereoptic depth, the children’s performance rapidly deteriorated. Even congenital cataract patients that were treated after the first 6 months of life demonstrated decreased performance in an illusory contour detection task (Putzar, Hötting, Rösler, & Röder, 2007), including also cases of monocular cataracts (Hadad, Maurer, & Lewis, 2017). Thus, it is evident that altered early-life experience resulting in visual deprivation can have lasting effects on mid-level visual functions such as contour completion and figure-ground segregation.

However, as our results show, such mid-level visual processes can be enhanced by the concurrent presentation of sounds. It is thus possible to imagine a multisensory training for children or young adults recovering from early visual deprivation in order to restore these mid-level visual functions. While practical limitations constitute an issue that too-often impedes the widespread use of multisensory technologies in clinical practice (Gori, Cappagli, Tonelli, Baud-Bovy, & Finocchietti, 2016), efforts are improving the accessibility of such treatment regimes, and are already demonstrating the utility of multisensory rehabilitation in visually deprived children (Cappagli, Finocchietti, Baud-Bovy, Cocchi, & Gori, 2017). Ongoing work from our group is introducing paradigms like that tested here into field research with sight-restored patients in rural India as well as into low-vision clinical units in Switzerland. The present study was a necessary proof-of-concept to validate

Another line of research investigating debilitating effects of early visual deprivation comes from studies examining auditory and haptic spatial impairments in early visually-deprived children (Cappagli et al., 2017; Gori et al., 2010), as well as impairments in multisensory processing in early visually-deprived adults (Champoux et al., 2010; Collignon, Charbonneau, Lassonde, & Lepore, 2009). Later in life some of these deficits might disappear, as early visually-deprived adults (for example through congenital blindness) demonstrate improved performance on various auditory spatial tasks (Collignon & De Volder, 2009; Collignon, Lassonde, Lepore, Bastien, & Veraart, 2007; Collignon, Renier, Bruyer, Tranduy, & Veraart, 2006; Collignon, Voss, Lassonde, & Lepore, 2009). Nevertheless, on some functions, such as auditory bisection tasks, early blind adults can still demonstrate lasting impairments (Gori, Sandini, Martinoli, & Burr, 2013). Thus, multisensory training programmes might pre-empt such cross-modal deficits in adults, while also improving multisensory integration in children that demonstrate such impairments.

There are instances where altered late-life experience, for example following brain injury, can lead to visual impairments (Dundon, Bertini, Làdavas, Sabel, & Gall, 2015). Neglect or hemianopic patients generally fail to report, respond to, or orient to visual stimuli presented contralaterally to the lesioned hemisphere (Halligan, Fink, Marshall, & Vallar, 2003), due to either a visual field deficit in hemianopia or to a visuospatial attentional deficit in neglect. Similarly, homonymous visual field defects are among the most serious deficits after cerebral artery stroke and traumatic brain injury in adults that result in either complete or partial loss of visual perception in one half of the visual field, and lead to numerous impairments in everyday functions (Dundon et al., 2015). In such patients, multisensory training improves not only their ocular functions, but also decreases their self-perceived disability in daily life activities such as bumping into objects, finding objects, and crossing the street (Bolognini, Rasi, Coccia, Ladavas, & Làdavas, 2005; Passamonti, Bertini, & Làdavas, 2009).

In conclusion, there is ample evidence of improvement of visual functions after multisensory training in both early and late visually-deprived individuals, and numerous open possibilities for rehabilitation or restoration of visual and multisensory functions. Our results add to this domain, by showing that mid-level vision can benefit from multisensory processes.

## Acknowledgements

Financial support for this work has been provided to M.M.M. by the Fondation Asile des aveugles (grant #232933), a grantor advised by Carigest SA (grant #232920), as well as the Swiss National Science Foundation (grant #169206).

## Author contributions

M.M. Murray developed the study concept. All authors contributed to the study design. Testing and data collection were performed by R.I. Tivadar. R.I. Tivadar and A. Gaglianese performed the data analysis and interpretation under the supervision of M.M. Murray. R.I. Tivadar drafted the manuscript, and A. Gaglianese and M.M. Murray provided critical revisions. All authors approved the final version of the manuscript for submission.

## Open Practices Statement

The data or materials for the experiments reported here are available upon reasonable request to the authors. None of the experiments was preregistered.

